# A novel cryo-electron microscopy support film based on 2D crystal of HFBI protein

**DOI:** 10.1101/2021.11.09.467987

**Authors:** Hongcheng Fan, Bo Wang, Yan Zhang, Yun Zhu, Bo Song, Haijin Xu, Yujia Zhai, Mingqiang Qiao, Fei Sun

## Abstract

Cryo-electron microscopy (cryo-EM) has become the most powerful tool to resolve the high-resolution structures of biomacromolecules in solution. However, the air-water interface induced preferred orientation, dissociation or denaturation of biomacromolecules during cryo-vitrification is still a major limitation factor for many specimens. To solve this bottleneck, we developed a new type of cryo-EM support film using the 2D crystal of hydrophobin I (HFBI) protein. The HFBI-film utilizes its hydrophilic side to adsorb protein particles via electrostatic interactions and keep air-water interface away, allowing thin-enough ice and high-quality data collection. The particle orientation distribution can be optimized by changing the buffer pH. We, for the first time, solved the cryo-EM structure of catalase (2.28-Å) that exhibited strong preferred orientation using conventional cryo-vitrification protocol. We further proved the HFBI-film is suitable to solve the high-resolution structures of small proteins including aldolase (150 kDa, 3.34-Å) and hemoglobin (64 kDa, 3.6-Å). Our work implied that the HFBI-film will have a wide application in the future to increase the successful rate and efficiency of cryo-EM.

## Introduction

With the recent breakthrough and developments, cryo-electron microscopy (cryo-EM) has become a powerful technique to study the high resolution three dimensional structures of biological macromolecules in solution^1^. Very recent reports also demonstrated that cryo-EM can even reach to atomic resolution^2,3^. However, there are still bottlenecks to overcome for a more successful rate and higher efficiency. One is how to routinely prepare the suitable cryo-vitrified samples for high resolution and high quality data collection^4^.

In the cryo-vitrification procedure using the conventional holey carbon support film, many challenges occurred, including protein denature/degradation and preferred orientation induced by the air-water interface^5^, non-uniform/ill distribution of protein particles in the hole^6^, un-expected few number of particles in the hole^7^, not well controlled ice thickness and etc., which have become an important barrier of high resolution structure determination for many biomacromolecules by cryo-EM.

Past efforts have been made to solve these challenges. One approach is changing the surface property of the holey carbon support film, e.g. by glow discharging^8^, special treatment with surfactants or PEG^9^. This approach can improve the particle distribution in the hole. Multiple blotting approach was tried to increase the number of particles in the hole^7^. Holy metal support film, including gold film^10,11^ and amorphous nickel titanium alloy film^12^, instead of carbon film, were also invented to improve the quality of the cryo-vitrified specimen. These approaches could not solve the air-water interface challenge.

Another approach is adjoining a layer of continuous film, e.g. ultra-thin carbon^13^, graphene^14–18^ or graphene derivate^19,20^ and 2D crystal of streptavidin^21,22^ et al., on the top of the holey carbon support film. This approach can increase the density of particles in the hole, protect the specimen away from the air-water interface, and alter the particle distribution. However, there exist non-negligible noisy background for ultra-thin amorphous carbon film, which limited the application for small proteins. In addition, the potential non-hydrophilic surface of carbon film or graphene materials would introduce a new factor to induce another preferred orientation of adsorbed particles.

Hydrophobins (HFBs) are a family of low molecular weight proteins (7~15 kDa) found in filamentous fungi^23^. HFBs contribute to the amphipathic film on the surface of fugal aerial hyphae and spores and play a key role in different physiological phases of fungi (Fig. 1a). HFBs exhibit a compact and globular structure that is stabilized by four conserved intramolecular disulfide bonds^24^. Interestingly, HFBs are amphiphilic proteins that display distinct hydrophobic and hydrophilic patches on the surface, which endows HFBs one family of the most surface-active proteins. HFBs can self-assemble into an amphipathic layer tightly adsorbed at the hydrophilic-hydrophobic interface such as the air-water interface or water-solid interfaces^25^. These unique properties of HFBs have led to many biotechnical applications including emulsions, drug delivery, tissue engineering^26^, biosensor^27,28^, protein purification/immobilization^29,30^ and so on. According to the hydropathy patterns and the properties of self-assembled films, HFBs are categorized into class I and class II^31,32^. The amphiphilic hydrophobin I protein (HFBI, 7.5 kDa) isolated from *Trichoderma reesei* is a kind of class II hydrophobins known to adhere to the air–water interface without denaturation and self-assemble into a crystalline mono-layer film (28 Å thickness)^33^. This self-assembled 2D crystal film at the air-water interface not only lowers the surface tension of the water but also has a very high surface shear elasticity that can stabilize air bubbles and foams^34^.

**Fig. 1.**
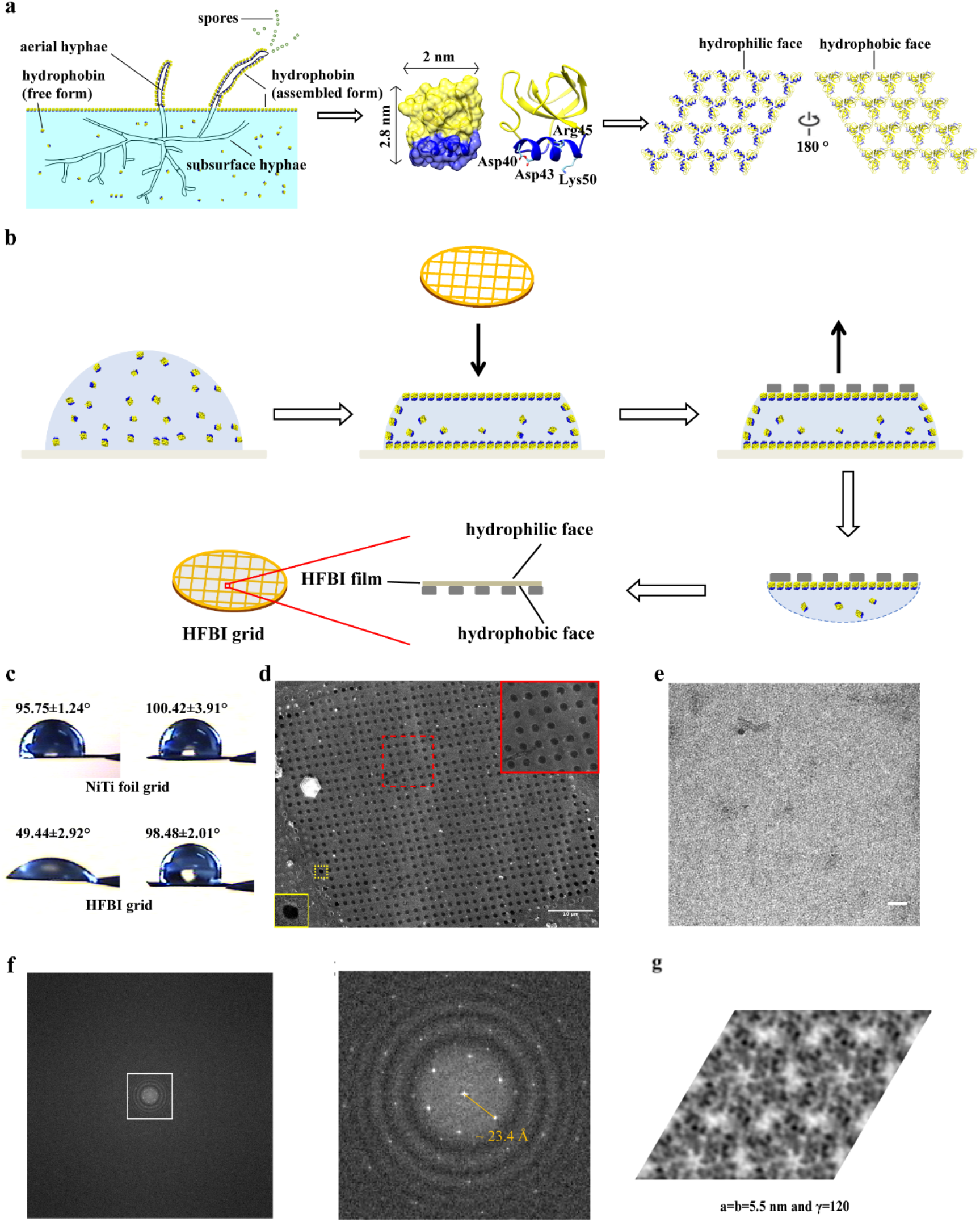
Production and characterization of HFBI-film covered cryo-EM grid. **a**, The biological nature of hydrophobin I protein (HFBI) and its physiological role during the filamentous fungal growth. HFBIs are secreted as monomers and assemble into 2D crystal film at the air-water interface, which coats the surfaces of aerial hyphae and spores and protects them from air-drying. **b**, The scheme of preparing HFBI-film covered cryo-EM grid. **c**, The water contact angles of NiTi foil grids without (upper) and with (bottom) HFBI-film coated were measured and labeled for both front (left) and back (right) sides. The HFBI-film was covered on the front side of the grid. No treatment of plasma cleaning was performed here. A drop of 3 μl water was used in this measurement. **d**, Cryo-SEM image of the HFBI-film covered on a holey NiTi foil grid. Most holes are covered by the HFBI-film. The regions in dotted squares are zoomed in at the insets, respectively. The region in the red square is completely covered by the HFBI-film and that in the yellow square contains am empty hole without the HFBI-film. Scale bar, 10 μm. **e**, The representative cryo-EM micrograph of the HFBI film. Scale bar, 20 nm. **f**, Power spectrum of the representative cryo-EM micrograph, showing the typical diffraction pattern of the HFBI 2D crystal with the first diffraction lattice spots around 23.4 Å. **g**, Imaging processing with unbending and Fourier extraction using 2dx package yields the arrangement of HFBI-film as the symmetry of p3. The unit cell parameter is labeled.

In the present study, we explored the potential application of HFBs to solve the air-water interface challenge in the cryo-EM field. We selected the 2D crystal of hydrophobin I as a new support film (HFBI-film) for cryo-vitrification procedure. We found that the HFBI-film is intrinsically hydrophilic with a relative low background and adsorbs protein particles via electrostatic interactions. By testing five different protein samples, apoferritin (480 kDa), glutamate dehydrogenase (340 kDa), catalase (240 kDa), aldolase (150 kDa) and hemoglobin (64 kDa), we proved that the HFBI-film can protect the specimen away from the air-water interface, solve the challenge of preferred orientation, improve the quality of cryo-EM micrograph by forming the thin ice, and be applicable to solve the high-resolution structure of small proteins with the molecular weight lower to 48 kDa.

## Results

### Preparation of cryo-EM grid covering by HFBI-film

We expressed recombinant HFBI in *Pichia pastoris* and got the protein purified by ultrafiltration and acetonitrile extraction (Extended Data Fig. 1a,b) according to our previous study^35^. Consistently, we found that HFBI can self-assemble into the well-ordered nanolayer (HFBI-film) with a thickness of 2 nm (Extended Data Fig. 1c,d). We also found HFBI nanolayer can alter the hydrophobic surface of siliconized glass to hydrophilic (Extended Data Fig. 1e,f) and alter the hydrophilic surface of mica to hydrophobic (Extended Data Fig. 1g,h) according to the water contact angle measurement, suggesting an amphipathic property of the HFBI-film (Fig. 1a).

We transferred the HFBI-film to a holey amorphous nickel titanium alloy (NiTi) foil grid^12^. After the visible and large HFBI-film spontaneously formed on the top of a solution drop, we incubated the NiTi foil grid on the top of the drop with the NiTi foil contacting the hydrophobic side of the HFBI-film, resulting the transfer of HFBI-film onto the grid (Fig. 1b). Since another exposed side of the HFBI-film is hydrophilic (Fig. 1c), the conventional grid treatments by plasma cleaning or ultraviolet (UV) irradiation before cryo-vitrification were not necessary for the HFBI-film. We then examined the integrity of HFBI-film by using cryo-scanning electron microscopy (cryo-SEM) and found that most area of the holey NiTi foil was covered by the HFBI-film uniformly and completely (Fig. 1d). In the subsequent cryo-EM experiment, we found that the HFBI-film had almost invisible background (Fig. 1e) with the power spectrum showing the first diffraction lattice spots around 23.4 Å (Fig. 1f). We then performed further image processing using the *2dx* software package^36^ and found that the lattice of the 2D crystal HFBI-film (Fig. 1g) is hexagonal with the symmetry of p3, which is in consistency with the previous study by atomic force electron microscopy^37^.

### HFBI-film allows thin enough ice and well-distributed particles

We first selected human apoferritin as the typical sample to explore the suitability of the HFBI-film for cryo-EM single particle analysis. We performed conventional cryo-vitrification procedure using FEI Vitrobot Mark IV by optimizing the parameter of blotting force and time. We found that, in comparison with the conventional cryo-EM grid, the grid covering the HFBI-film can enable a uniform much thinner ice containing apoferritin particles (Fig. 2a,b). We took serials of electron micrographs using various defocuses from −1.92 μm to −0.48 μm (Extended Data Fig. 2) and found that the apoferritin particles showed a nice contrast even at a small defocus value of −0.48 μm (Fig. 2a), implying that the ice was thin enough. We also found that the apoferritin particles showed a uniform dispersed and low-aggregated distribution, suggesting a high quality of cryo-vitrified specimen. Our subsequent cryo-electron tomographic reconstruction indicated that almost the particles within the ice formed a single thin layer and adsorbed onto the HFBI-film, avoiding the air-water interface contact (Extended Data Fig. 3).

**Fig. 2.**
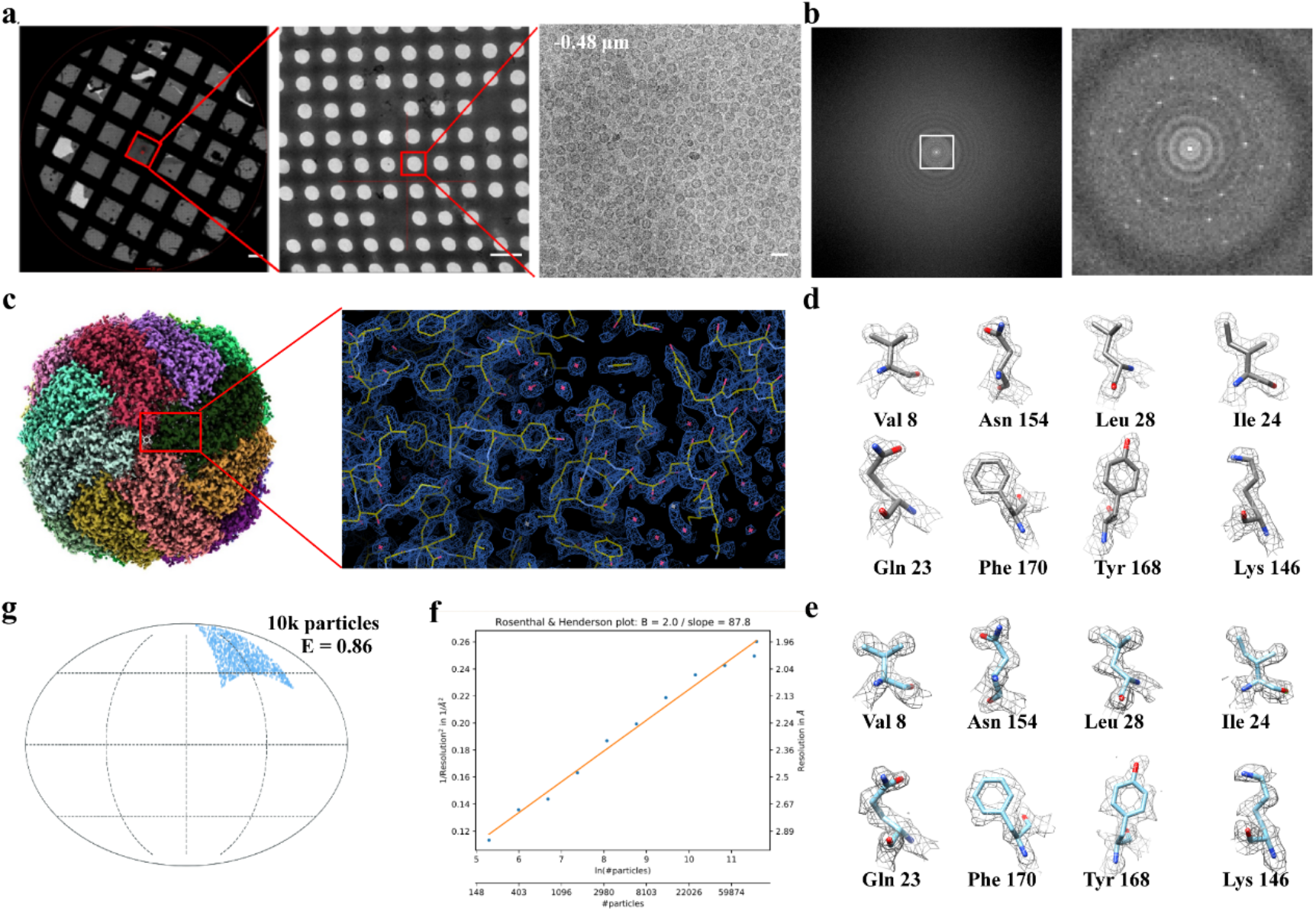
Cryo-EM application of HFBI-film using human apoferritin as a test sample. **a**, Cryo-EM micrographs of cryo-vitrified grid in different magnifications showing uniform and thin vitrified ice as well as the uniform distribution of apoferritin particles, respectively. The defocus value of the micrograph in high magnification (right) is labeled. Scale bar, from left to right, 20 μm, 2 μm, 20 nm. **b**, Power spectrum (left) of the high magnification cryo-EM micrograph in **a**. The central region is zoomed in (right) showing the diffraction spots of 2D crystal of HFBI-film. **c**, Cryo-EM map of the human apoferritin at 1.96-Å resolution and its representative region is zoomed in (right) with atomic model fitted. **d**, Representative electron densities of selected amino acid residues. **e**, Representative electron densities of selected amino acid residues after density modification. **f**, Rosenthal-Henderson plot with the estimated B-factor of 87.8 Å. **g**, The orientation distribution of apoferritin particles with the cryo-EF value of 0.86 calculated using 10,000 particles.

Using this HFBI-film covered cryo-EM grid, we collected the cryo-EM dataset of human apoferritin by using FEI Talos Arctica operated in 200 kV and equipped with GATAN BioQuantum K2 camera (Extended Data Table 1). After several steps of single particle analysis (Extended Data Fig. 4a,b), we obtained the final 3D cryo-EM map of human apoferritin (Fig. 2c) at an average resolution of 1.96-Å (Extended Data Fig. 4c) according to the gold standard Fourier Shell Correlation threshold FSC_0.143_ (Extended Data Fig. 4d). At the resolution of 1.96-Å, we could clearly see the holes in aromatic residues such as phenylalanine and tyrosine (Fig. 2d). We further performed density modification^38^ to improve the quality of the map (Fig. 2e), which is comparable to the recently reported cryo-EM map of apoferritin at a resolution of 1.75-Å, the highest resolution reached by using 200 kV cryo-electron microscope^39^. To our best knowledge, our work presented the highest resolution of cryo-EM single particle analysis that used the continuous thin support film covered grids, including the graphene monolayer covered grids^16,18^.

We estimated the Rosenthal-Henderson B-factor^40^ of this dataset as 87.8 Å^2^ (Fig. 2f), suggesting that we could only use 200~400 particles of this dataset to reach sub-3 Å resolution. We noted that the B-factor of the 1.75-Å apoferritin map is 90.7 Å^239^. We believe it was the thin enough ice of the cryo-vitrified specimen to yield the high quality of the dataset with the small B-factor. We further analyzed the apoferritin particle orientation distribution in our dataset and found an even distribution (Extended Data Fig. 4e) with the calculated cryo-FE^41^ value of 0.86 (Fig. 2g), which is much better than that of the previously reported apoferritin cryo-EM dataset using the graphene monolayer covered grids^16^.

### HFBI-film solves the strong preferred orientation of catalase

It has been shown that the air-water interface using the conventional cryo-EM grid would induce 90% particles attaching to the air-water interface, resulting a significant preferential orientation of particles in many cases^42^. Our above study of the HFBI-film using the human apoferritin gave the insight that the HFBI-film could protect the protein particles away from the air-water interface and thus avoid the potential preferential orientation of particles. We then selected catalase as another typical sample to prove (Extended Data Table 1).

Using the conventional cryo-EM sample preparation protocol, catalase exhibited a strong preferred orientation with its side view^43^ (see also Fig. 3a,b), which has forbidden us to use cryo-EM single particle analysis to solve the high-resolution structure of catalase. However, after cryo-vitrification of catalase using the HFBI-film covered NiTi foil grid, we observed a uniform distribution of catalase particles with various views, which was further assessed by 2D classification (Fig. 3c,d). After several steps of single particle analysis (Extended Data Fig. 5a), we obtained the cryo-EM map of catalase at the resolution of 2.28-Å (Fig. 3e,f and Extended Data Fig. 5b) according to the gold standard threshold FSC_0.143_ (Extended Data Fig. 5c). This is the first time for us to resolve the near atomic resolution structure of catalase using cryo-EM single particle analysis approach.

**Fig. 3.**
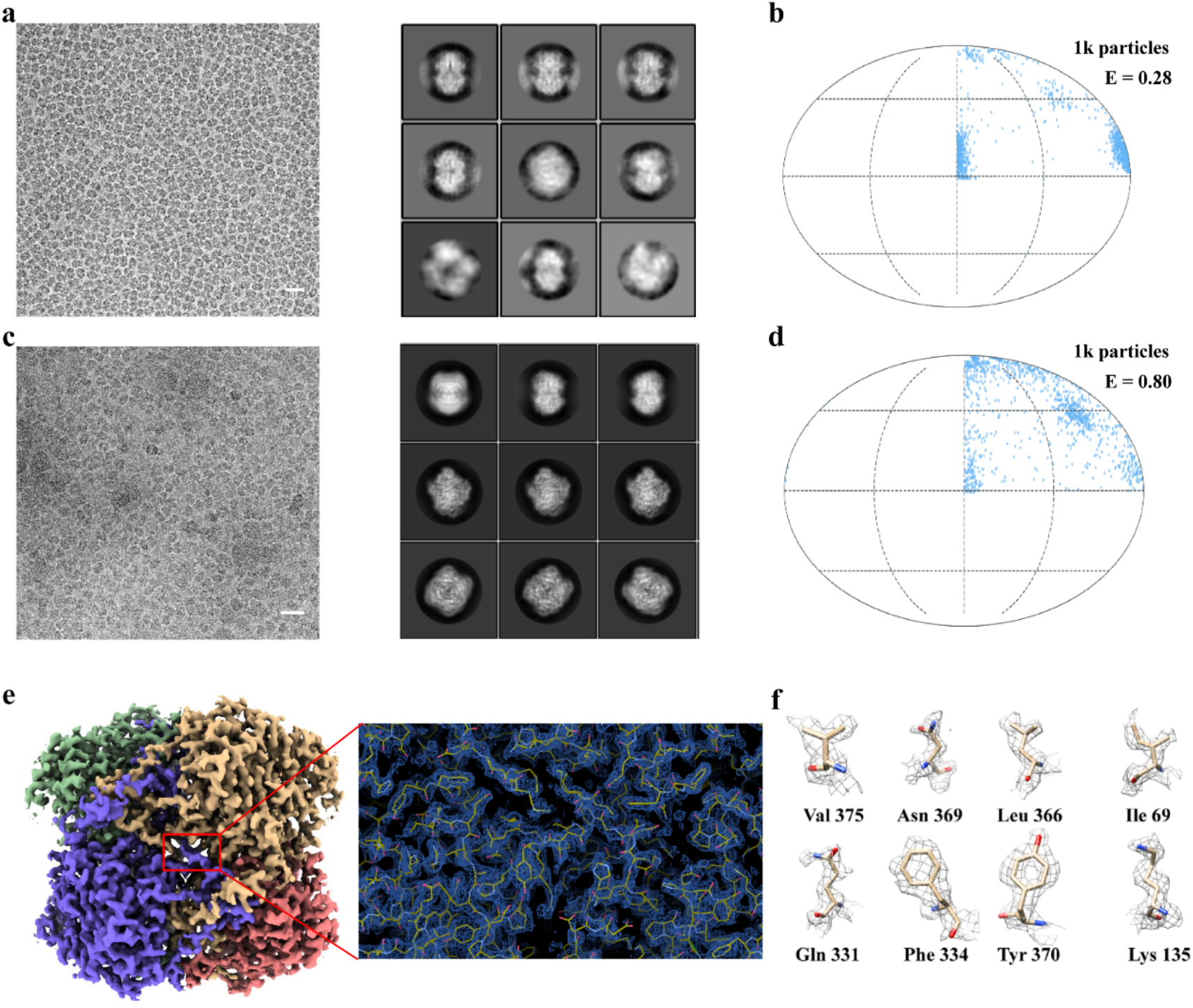
Using HFBI-film to solve the preferred orientation problem. **a**, A representative cryo-EM micrograph of catalase using a holey NiTi foil grid (scale bar, 20 nm) and the representative 2D class averages. **b**, The orientation distribution of catalase particles with the cryo-EF value of 0.28 calculated using 1,000 particles in the dataset of **a**. **c**, A representative cryo-EM micrograph of catalase using HFBI-film covered NiTi foil grid (scale bar, 20 nm.) and the representative 2D class averages. **d**, The orientation distribution of catalase particles with the cryo-EF value of 0.80 calculated using 1,000 particles in the dataset of **b**. **e**, Cryo-EM map of the catalase at 2.28-Å resolution and its representative region is zoomed in (right) with atomic model fitted. **f**, Representative electron densities of selected amino acid residues.

We further analyzed the particle orientation distribution in the dataset using the HBFI-film and observed an even distribution of catalase particles (Extended Data Fig. 5f) with the calculated cryo-EF value of 0.80 (Fig. 3d and Extended Data Fig. 5e), which is significantly improved in comparison with the cryo-EF value of 0.28 in the dataset using the NiTi foil grid only (Fig. 3b). We noted that the cryo-EF value of the previous study using the holey pure gold foil grid was reported as 0.2^43^. As a result, the HBFI-film can solve the preferred orientation problem by preventing particles from interacting with the air-water interface, thus enabling structure determination to the near-atomic resolution.

In addition, similar to the above study of human apoferritin, since the thickness of the ice could be thin enough by using the HFBI-film, we could collect a high quality of cryo-EM dataset of catalase with a small Rosenthal-Henderson B-factor of 79.3 Å^2^ (Extended Data Fig. 5d).

### pH regulated particle orientation distribution on HFBI-film

Considering the hydrophilic nature of the exposed side of HFBI-film, we believe the protein particles are adsorbed mainly by electrostatic interaction^44^. To prove this hypothesis, we selected glutamate dehydrogenase (GDH) as a third typical sample to study its orientation distribution in the cryo-vitrified sample using the HFBI-film and study whether such distribution could be regulated by the pH value (Extended Data Table 1).

We found that the orientation distributions of GDH were obviously different upon different pH conditions (Fig. 4a and Extended Data Fig. 6). In the acidic condition GDH particles majorly showed top and oblique views, while in the basic condition GDH mainly showed side views (Fig. 4b). We collected a large cryo-EM dataset of GDH at the pH value of 7.5, which is close to its isoelectric point (pH 7.66). At this pH condition, GDH particles exhibited a most even distribution. After several steps of single particle analysis (Extended Data Fig. 7a), we obtained the cryo-EM map of GDH at the resolution of 2.28-Å (Fig. 4c and Extended Data Fig. 7c) according to the gold standard threshold FSC_0.143_ (Extended Data Fig. 7d). The cryo-EF analysis of this dataset gave the factor of 0.84 (Extended Data Fig. 7e) and indicated there is no preferred orientation issue (Extended Data Fig. 7f). We also calculated the Rosenthal-Henderson B-factor of this dataset as 76.0 Å^2^ (Extended Data Fig. 7g), again suggesting a high quality of cryoEM dataset.

**Fig. 4.**
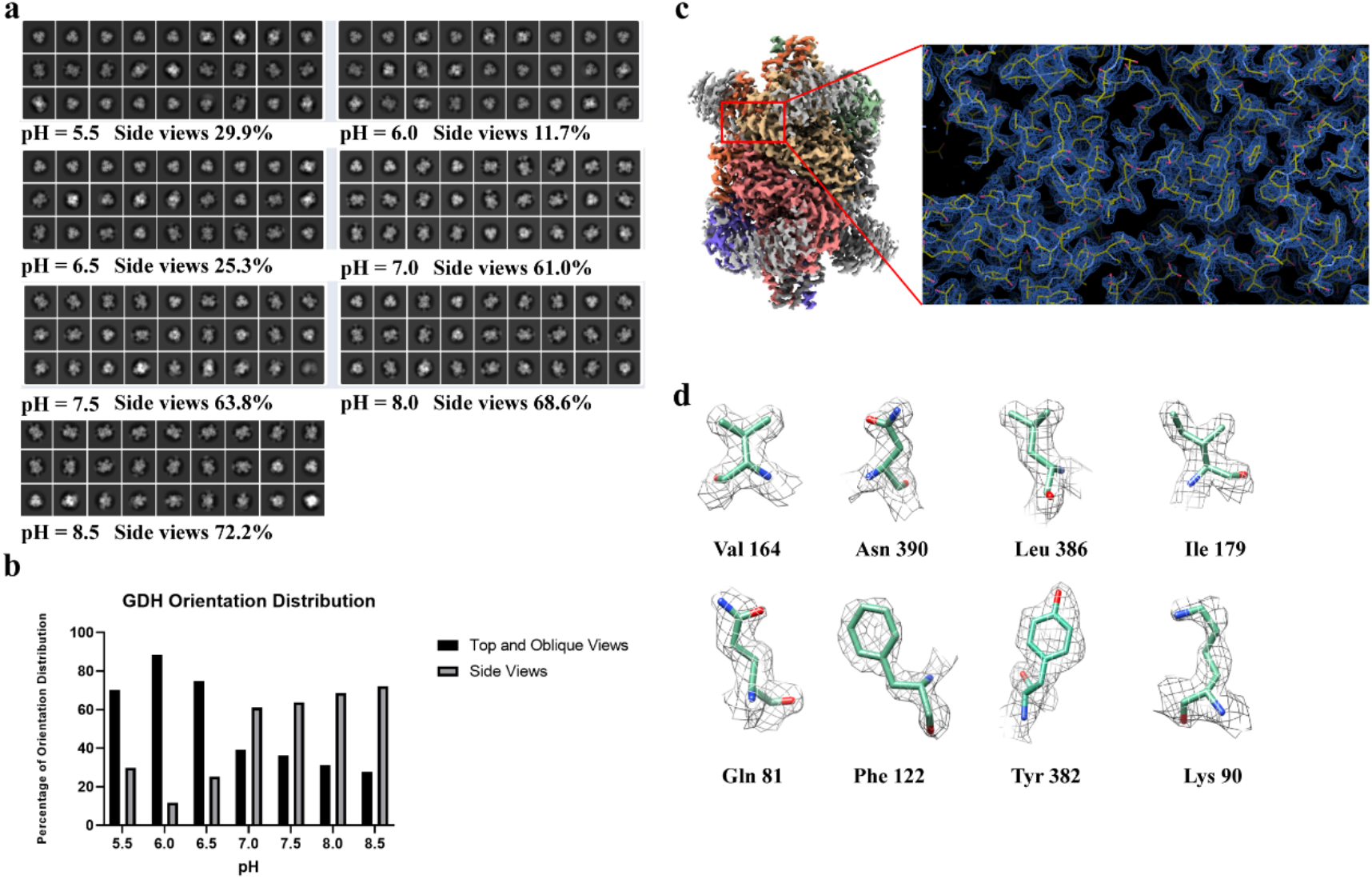
Regulating particle orientation distribution by adjusting the pH. **a**, Representative 2D class averages of GDH in different pH values using the HFBI-film covered grid. The percentages of side views of GDH were counted and labeled accordingly. **b**, The statistics of percentages of different GDH views at different pH values. **c**, Cryo-EM map of GDH in pH 7.5 was solved to 2.28-Å resolution and its representative region is zoomed in (right) with atomic model fitted. **d**, Representative electron densities of selected amino acid residues.

As a result, although the HFBI-film could raise a new preferential orientation distribution like other thin continuous support films, e.g. graphene or ultra-thin carbon film, the particular electrostatic adsorption mechanism of the HFBI-film could allow us to optimize the distribution of particles by adjusting the pH value. In principle, we believe, the pH value could be optimized to the isoelectric point of the target protein.

### Application of HFBI-film for small protein particles

To further explore the potential of our HFBI-film for cryo-EM experiments of small proteins (<200 kDa), we selected aldolase (150 kDa, homo-tetramer) and hemoglobin (64 kDa, hetero-tetramer αβαβ) as the test samples (Extended Data Table 1).

With the thin enough ice using the HFBI-film, we could clearly visualize the particles of aldolase and hemoglobin from raw electron micrographs (Fig. 5a,d) and could observe the secondary elements directly from the 2D classifications (Extended Data Fig. 8b,10b). After several steps of single particle analysis (Extended Data Fig. 8a), we obtained the cryo-EM map of aldolase at the resolution of 3.34-Å (Fig. 5b,c and Extended Data Fig. 8c) according to the gold standard threshold FSC_0.143_ (Extended Data Fig. 8d). We did not observe any preferential orientation with the computed cryo-EF value of 0.84 (Extended Data Fig. 8e,f). We found that the calculated Rosenthal-Henderson B-factor (156.3 Å^2^, Extended Data Fig. 8g) of the cryo-EM dataset of aldolase was double larger than those of the above studies for apoferritin, catalase and GDH.

**Fig. 5.**
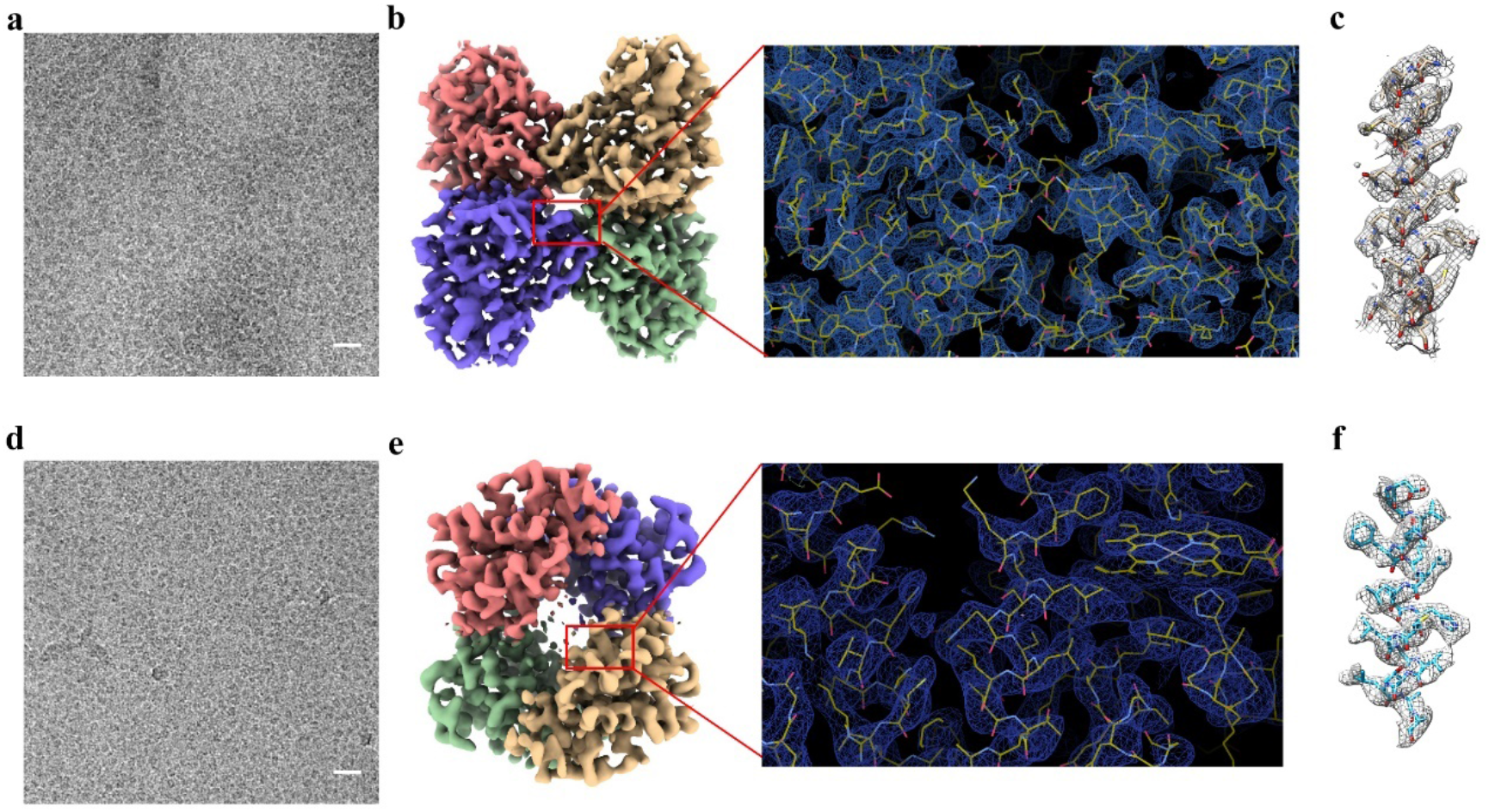
Applying HFBI-film to small proteins. **a**, The representative cryo-EM micrograph of aldolase. Scale bar, 20 nm. **b**, Cryo-EM map of the aldolase at 3.34-Å resolution and its representative region is zoomed in (right) with the atomic model fitted. **c**, Representative map of one helix of the aldolase with the atomic model fitted. **d**, The representative cryo-EM micrograph of hemoglobin. Scale bar, 20 nm. **e**, Cryo-EM map of the hemoglobin at 3.6-Å resolution and its representative region is zoomed in (right) with the atomic model fitted. **f**, Representative map of one helix of the hemoglobin with the atomic model fitted.

The increased Rosenthal-Henderson B-factor of aldolase dataset would be due to the background of the HFBI-film. Therefore, to process the cryo-EM dataset of hemoglobin with the molecular weight of 64 kDa, we began to consider the significant influence of this factor. The signal of the 2D crystal HFBI-film could be simply removed using a lattice filtering algorithm in Fourier space^45,46^ (Extended Data Fig. 9). Starting from the background extracted dataset, after several steps of single particle analysis (Extended Data Fig. 10a), we obtained the cryo-EM map of hemoglobin at the resolution of 3.6-Å (Fig. 5e,f and Extended Data Fig. 10c) according to the gold standard threshold FSC_0.143_ (Extended Data Fig. 10d). No orientation bias was found with the cryo-EF values of 0.86 (Extended Data Fig. 10e,f). The calculated Rosenthal-Henderson B-factor (162.2 Å^2^, Extended Data Fig. 10g) was again double larger than those of the above studies for apoferritin, catalase and GDH.

We further explored whether our HFBI-film could be suitable for even smaller proteins. Utilizing a similar approach of the previous study^17^, starting from the raw dataset of 64 kDa hemoglobin particles, we performed *in silica* substation (Extended Data Fig. 11) to generate two datasets of smaller particles, one for a heterodimer (αβ subunit, 32 kDa), and another for a heterotrimer (αβα subunit, 48 kDa). For each dataset, we performed the reference-free 2D classification and the 2D classification with pre-aligned angular information (Extended Data Fig. 11a). For the dataset of the heterotrimer, we found that the reference-free classification with the global alignment could still yield good class averages with the secondary structural details. However, for the dataset of the heterodimer, we observed many classes with mis-aligned particles, comparing to the classification using the prior alignment. Consistently, our subsequent 3D reconstructions by performing angular global search from scratch also showed that the 48 kDa heterotrimer could be resolved successfully to a high resolution of 3.8-Å while the 32 kDa heterodimer could only be resolved to a low resolution of 6.4-Å (Extended Data Fig. 11b,c), suggesting the signal-to-noise ratio of the 32 kDa heterodimer particles at the current dataset was not enough for precise particle alignment.

As a result, our above studies proved that the HFBI-film could be applied to study the high resolution cryo-EM structures of small proteins with the molecular weight lower to 48 kDa.

## Discussion

In the present study, we developed a new type of cryo-EM grid by coating 2D crystal of HFBI protein onto the holey amorphous NiTi foil grid and evaluated this kind of grid to solve the air-water interface problem, which has been recognized as the major factor to induce denaturation/aggregation and preferred orientation of protein particles during the conventional vitrification procedure and prevent the high resolution study of cryoEM

The amphipathic property of the HFBI-film allows it easy to coat onto the holey amorphous NiTi foil using its hydrophobic side and leave its hydrophilic side exposed ready to adsorb the protein particles via electrostatic interactions. Thus, unlike other cryo-EM grids, pre-treatment such as plasma cleaning or UV irradiation before cryo-vitrification is not necessary. The adsorption of protein particles by the HFBI-film not only protects the specimen away from the air-water interface but also allows the thickness of the ice thin enough, which is important to increase the quality of the cryoEM images, which has been proved by the small Rosenthal-Henderson B-factor in our experiments. With the HFBI-film, we for the first time determined the high resolution cryo-EM structure of catalase, which showed a severe air-water interface induced preferential orientation using the conventional cryo-vitrification protocol.

Although the preferential orientation issue also exists by using HFBI-film, the orientation distribution of protein particles can be optimized by adjusting the pH of the buffer to regulate the interaction between the specimen and the HFBI-film. We speculated that the optimal pH could be selected close to the isoelectric point of the target protein. Thus, the potential preferential orientation issue by using HFBI-film can be easily solved by adjusting the pH value, which has been proved by our experiments using GDH as a test sample.

The thickness of the HFBI-film is 2 nm, which is thin enough to keep a low background for most cryo-EM experiments. In our present study, we proved that a standard image processing procedure was enough for those particles with the molecular weight higher than 150 kDa. With the lower molecular weight, the Rosenthal-Henderson B-factor increased and the achievable resolution decreased (Extended Data Fig. 12). For those protein particles with the molecular weight lower than 100 kDa, additional background removing procedure would be important for high resolution reconstruction. Since the HFBI-film is the 2D crystal, its lattice background can be easily removed by Fourier filtering algorithm without damaging the quality of the data. We proved that with this procedure the structure of human hemoglobin (64 kDa) can be successfully solved to 3.6-Å. By additional *in silico* experiments, we further showed that at least a protein particle with the molecular weight of 48 kDa could be solved to the high resolution by using our HFBI-film.

Compared to other types of continuous support films such as graphene and 2D crystal of streptavidin, the production of our HFBI-film is simple and efficient, the pre-treatment before cryo-vitrification is not needed, the background is low and suitable for small particles, and the particle adsorption mechanism is new via electrostatic interactions, allowing regulating particle orientation distribution simply by changing the pH value. Thus, the use of our HFBI-film is an economic and highly efficient solution with a high successful rate to solve the air-water interface challenge.

Overall, our present work proves a new type of continuous support film coated grid that would have a wide application in the future cryo-EM studies.

## Methods

### Expression and Purification of Hydrophobin I protein

Molecular cloning and recombinant expression of HFBI have been reported previously^35^. Briefly, positive transformants (His+ Mut+) were chosen for flask cultivation and induced in buffered minimal methanol (BMM) medium with 0.7% methanol at 30 °C for 4 days. One strain with the highest production of HFBI was chosen for 96 h Fed-batch fermentation.

HFBI was purified by a two-step procedure including ultrafiltration and acetonitrile extraction. The expression supernatant containing HFBI was collected and purified by ultrafiltration using a hollow fiber membrane module (Tianjin MOTIMO Membrane Technology Ltd., China) and then lyophilized into powders. The powders were redissolved in 0.01% trifluoroacetic acid / 40% acetonitrile solution with a final concentration of 40 mg ml^-1^, stirring for 10 min prior to ultrasonic for 10 min. The crude extracted proteins were collected by the centrifugation with 8000 rpm for 30 min and then lyophilized into powders and stored for further study.

### Water contact angle (WCA) measurements

Hydrophobic siliconized glass (HR3-239; Hampton Research) and hydrophilic mica sheets (J & K Chemical Technology, Tianjin, China) were used to study the properties of HFBI-film on solid surfaces. Each surface was coated with 50 μl of 100 μg ml^-1^ HFBI solution and incubated at room temperature (25 °C) overnight, using uncoated substrates as controls. The surface was subsequently rinsed three times with Milli-Q water to remove uncoated proteins and dried naturally. WCA was measured with a KSV Contact Angle Measurement System with a 5 μl water droplet on the protein-coated side of the material. Three replicates were performed on different areas of the sample surface.

### Atomic force microscopy (AFM) measurements

The freshly cleaved mica was coated with a 100 μl droplet of purified HFBI protein solution (100 μg ml^-1^) and incubated for 3 min at room temperature, rinsed three times with Milli-Q water to remove uncoated proteins and dried naturally. The topography of HFBI-coated mica surface was investigated by atomic force microscopy (AFM, Dimension Icon, Bruker, Frankfurt, America). The AFM parameters were set as follows, scan size of 5 μm, scan rate of 1.95 Hz, resolution of 512×512 pixels, and aspect ratio of 1.0. The AFM images were processed using the NanoScope Analysis software 1.5.

### Preparation of HFBI-film covered grid

The purified and lyophilized powder of HFBI protein was dissolved in milliQ water (pH 7) with a concentration of 3.6 mg ml^-1^, followed by sonication for about 1min to prevent from forming aggregation. Then the solution was further diluted to the working concentration of 100 μg ml^-1^, followed by additional sonication. A 30 μl drop of the working solution was added onto the surface of parafilm that was placed in a closed cell-culture dish and incubated for ~ 3 h in a humidified environment at room temperature and atmospheric pressure. When a visible film formed on the top of the drop, we transferred the film to a 300-mesh amorphous NiTi foil 1.2/1.3 Au grid (Zhenjiang Lehua Electronic Technology, China) by incubating the grid on top of the drop for 15 min with the NiTi foil film in contact with the HFBI-film (Fig. 2b). After the transfer, a 5 μl drop of milliQ water was added twice on the HFBI-film to clean off the impurities and a filter paper was used to draw remaining water away from the grid. Then the grid was left to dry in a grid holder at room temperature for the subsequent cryo-EM experiments.

### Preparation of tested sample and cryo-vitrification

The human apoferritin was provided by Prof. K.L. Fan and Prof. X.Y. Yan’s lab (Institute of Biophysics, CAS) and diluted to a concentration of 2 mg ml^-1^ in the buffer containing 20 mM Tris-HCl and 150 mM NaCl (pH 6.5). The catalase from human erythrocytes, GDH, lyophilized rabbit muscle aldolase and lyophilized human hemoglobin were purchased from Sigma Aldrich. The catalase was diluted to a final concentration of 2.3 mg ml^-1^ in the buffer of PBS (pH 6.5). The GDH was diluted to a final concentration of 3 mg ml^-1^ and exchanged into the buffer of PBS (pH 7.5) by ultra-filtration. Lyophilized rabbit muscle aldolase was solubilized in the buffer of 20mM HEPES pH 7.5, 50 mM NaCl to a final concentration of 1.2 mg ml^-1^. Lyophilized human hemoglobin was solubilized in the buffer of PBS (pH 7.5) to a final concentration of 6.3 mg ml^-1^. All these specimens were used directly for the subsequent cryo-vitrification without further treatments.

Using Vitrobot Mark IV (ThermoFisher Scientific, USA), 3.0 μl aliquots of the sample were applied to the normal NiTi grid or the HFBI-film coated grid at 4 °C and 100 % humidity. After waiting for 2 min, the grid was blotted for about 4 s and rapidly plunged into the liquid ethane for cryo-vitrification.

### Cryo-EM data collection

The cryo-EM dataset of apoferritin was collected in an FEI Talos Arctica electron microscope (200 kV) equipped with an energy filter and direct electron detector (Gatan BioQuantum K2) operated at the super resolution mode (**Extended Data Table 1**). A total number of 2649 raw movie stacks were automatically collected using SerialEM^47^. Images were recorded by beam-image shift method^48^ with a physical pixel size of 0.8 Å. The total dose was 50 e^-^/Å^2^ to generate 32-frame gain normalized stacks in TIFF format. The defocus range varied from −0.6 μm to −2 μm.

The dataset of catalase was collected in an FEI Titan Krios G2 electron microscope (300 kV) equipped with an energy filter and direct electron detector (Gatan BioQuantum K2) operated at the super resolution mode (**Extended Data Table 1**). For the dataset using the HFBI-film coated grids, a total number of 372 movies were automatically collected by SerialEM with a physical pixel size of 0.82 Å. For the dataset using the normal NiTi grids, a total number of 73 movies were automatically collected using SerialEM with a physical pixel size of 0.82 Å. To push high resolution, another large dataset using the HFBI-film coated grids was collected with 2842 movies and a physical pixel size of 0.65 Å. Images were recorded by beam-image shift method. The total dose was 60 e^-^/Å^2^ to generate 40-frame gain normalized stacks in TIFF format. The defocus range varied from −0.6 μm to −1.6 μm.

The dataset of GDH was collected in an FEI Titan Krios G2 electron microscope (300 kV) equipped with an energy filter and direct electron detector (Gatan BioQuantum K2) operated at the super resolution mode (**Extended Data Table 1**). A total number of 1635 raw movie stacks were automatically collected by SerialEM. Images were recorded by beam-image shift method with a physical pixel size of 0.65 Å. The total dose was 60 e^-^/Å^2^ to generate 40-frame gain normalized stacks in TIFF format. The defocus range varied from −0.6 μm to −1.6 μm.

The dataset of aldolase was collected in an FEI Titan Krios G2 electron microscope (300 kV) equipped with an energy filter and direct electron detector (Gatan BioQuantum K2) operated at the super resolution mode (**Extended Data Table 1**). A total number of 2460 raw movie stacks were automatically collected by SerialEM. Images were recorded by beam-image shift method with a physical pixel size of 0.65 Å. The total dose was 70 e^-^/Å^2^ to generate 40-frame gain normalized stacks in TIFF format. The defocus range varied from −0.6 μm to −1.6 μm.

The dataset of human hemoglobin was collected in an FEI Talos Arctica electron microscope (200 kV) equipped with an energy filter and direct electron detector (Gatan BioQuantum K2) operated at the super resolution mode (**Extended Data Table 1**). A total number of 2766 raw movie stacks were automatically collected by SerialEM. Images were recorded by beam-image shift method with a physical pixel size of 0.65 Å. The total dose was 70 e^-^/Å^2^ to generate 45-frame gain normalized stacks in TIFF format. The defocus range varied from −0.6 μm to −1.6 μm.

### Cryo-EM data processing

All movie stacks were subjected to beam-induced motion correction and dose-weighting using RELION3.1 with a 5 x 5 patch and a 2-fold binning^49,50^. Contrast transfer function parameters for each micrograph were estimated by Gctf1.06^51^. Protein particles from a small subset of micrographs were picked by Gautomatch (developed by Kai Zhang, http://www.mrc-lmb.cam.ac.uk/kzhang/Gautomatch/). These particles were subjected into multiple rounds of 2D classification to generate clean particles for the subsequent topaz training^52^. The topaz method was used for the automatic particle picking based on the deep-denoised micrographs^53^. In total, there were 351,565 particles for the apoferritin, 653,922 particles for the catalase, 35,587 particles for the glutamate dehydrogenase, 769,925 particles for the aldolase and 874,836 particles for the hemoglobin picked, respectively. Then, subsequent multiple rounds of 2D classification and 3D classification of binned particles in RELION3.1 were performed to discard “junk” particles.

For the datasets of apoferritin, glutamate dehydrogenase, aldolase and catalase, 3D auto-refinement, CTF Refinement and Bayesian polishing were all performed in RELION3.1 using the standard procedure. RELION ’s local resolution estimation was used to calculate the local resolution map. Final maps were all post-processed in RELION for model building and refinement. Density modification of apoferritin cryoEM map was performed in PHENIX (v1.19-4092)^54^ using the module of ResolveCryoEM^38^.

For the dataset of hemoglobin, the stander processing procedures including 2D classification and 3D classification were first performed in RELION. Then, the background of the HFBI 2D crystal lattice was removed from the motion-corrected micrographs using a modified Fourier filtering scripts^45,46^. The particles were then re-centered and re-extracted from the cleaned micrographs in RELION with the original pixel size of 1.04 Å. The subsequent 3D auto-refinement with C2 symmetry and post-processing yielded a map with a global resolution of 4.25 Å. Then the refined particles were subjected to an additional 3D classification in the local search mode. 103,777 good particles were selected and subjected to 3D auto-refinement with C2 symmetry enforced. The global resolution was improved to 4.0 Å. The particles were then subjected to Bayesian polishing, pushing the resolution to 3.9 Å. We then performed one round of random phase 3D classification^55^ to further remove bad particles. The final 58,828 good particles were subjected for further 3D auto-refinement and Bayesian polishing, pushing the resolution to 3.69 Å. Finally, we imported the RELION refined particles into M^56^ and improved the final resolution to 3.6 Å for the subsequent model building and refinement. Here, M^56^ was also used to calculate the local resolution map.

### Model fitting and refinement

The atomic models of apoferritin (PDB code 7K3V), catalase (PDB code 1DGH), glutamate dehydrogenase (PDB code 5K12), aldolase (PDB code 6V20) and hemoglobin (PDB code 5NI1) were docked into the corresponding cryo-EM maps using UCSF-Chimera^57^. Then each model was manually adjusted in Coot^58^ and refined in real space using PHENIX^54^ (**Extended Data Table 1**).

### Cryo electron tomography data acquisition and processing

We collected cryo electron tomography dataset of the same apoferritin that was used for cryo-EM single particle data collection. Tilt series were collected from −54° to 54° with a 3° interval at ×79,000 magnification (EFTEM mode, 1.7 Å pixel size) in an FEI Talos Arctica electron microscope (200 kV) equipped with an energy filter and direct electron detector (Gatan BioQuantum K2). For each tilt, the exposure time was set 1.0 s with 8 frames using a total dose of 3.38 e-/Å^2^ in electron counting mode. Each set of tilt series has a total dose of 125 e-/Å^2^. Data acquisition was automatically performed using SerialEM^47^. The tilt series raw stacks were motion-corrected by MotionCor2^59^. The fiducial-free alignment and tomogram reconstruction were processed in EMAN2.3^60^. The final tomograms were generated with four-time binning (pixel size 8.4 Å).

## Data availability

The atomic coordinates and cryo-EM density maps of apoferritin, catalase, glutamate dehydrogenase, HA trimer, aldolase and haemoglobin have been deposited in the RCSB Protein Data Bank (PDB) with the accession codes 7VD8, 7VD9, 7VDA, 7VDF, 7VDC and 7VDE and in the Electron Microscopy Data Bank (EMDB) with the accession codes EMD-31910, EMD-31911, EMD-31912, EMD-31916, EMD-31913 and EMD-31915, respectively.

## Code availability

The scripts used for Fourier filtering are available at the accompanying Mendeley data site http://dx.doi.org/10.17632/k2g2p5z9x6.2. We further modified the Fourier filtering scripts according to the HFBI crystal lattice parameters.

## Acknowledgements

We are grateful to Ping Shan, Ruigang Su and Mengyue Lou (F.S. lab) for their assistance in the lab management. All the sample preparation and cryoEM works were performed at Center for Biological Imaging (CBI, http://cbi.ibp.ac.cn), Institute of Biophysics, Chinese Academy of Sciences. We would like to thank B.Z, X.H., X.L., T.N. and L.C. from CBI for their help with the cryo-EM data collection. We would be grateful to X.J. from CBI for her help in cryo-SEM experiments. This work was equally supported by grants from the Ministry of Science and Technology of China (2017YFA0504700), National Natural Science Foundation of China (31830020), and Chinese Academy of Sciences (XDB37040102). The authors would also like to thank the grant supports from National Natural Science Foundation of China for Distinguished Young Scholars (31925026), from Beijing Municipal Science and Technology Commission (Z181100004218002) and the Sino-Swiss scientific and technological cooperation project by the Ministry of Science and Technology of China (2015DFG32140).

## Author contributions

F.S. initiated the project. F.S. and M.Q supervised the project. H.F. designed and performed the production of HFBI-film covered grids. B.W. expressed, purified and characterized HFBI proteins. H.F. performed all the cryo-EM sample preparation, data collection, imaging processing and model building. H.F., F.S., Yan Z., Yun Z. and Yujia Z. analyzed the data. H.F. and F.S. wrote the manuscript with additional input from B.W. and M.Q..

## Competing interests

Parts of this study (the production of HFBI-film covered grid) has been submitted as a Chinese patent of invention with the application number of 202110576212.4 and is currently under scrutiny.

## Correspondence and requests for materials

should be addressed to F.S. or M.Q.

---

**Extended Data Table 1.**
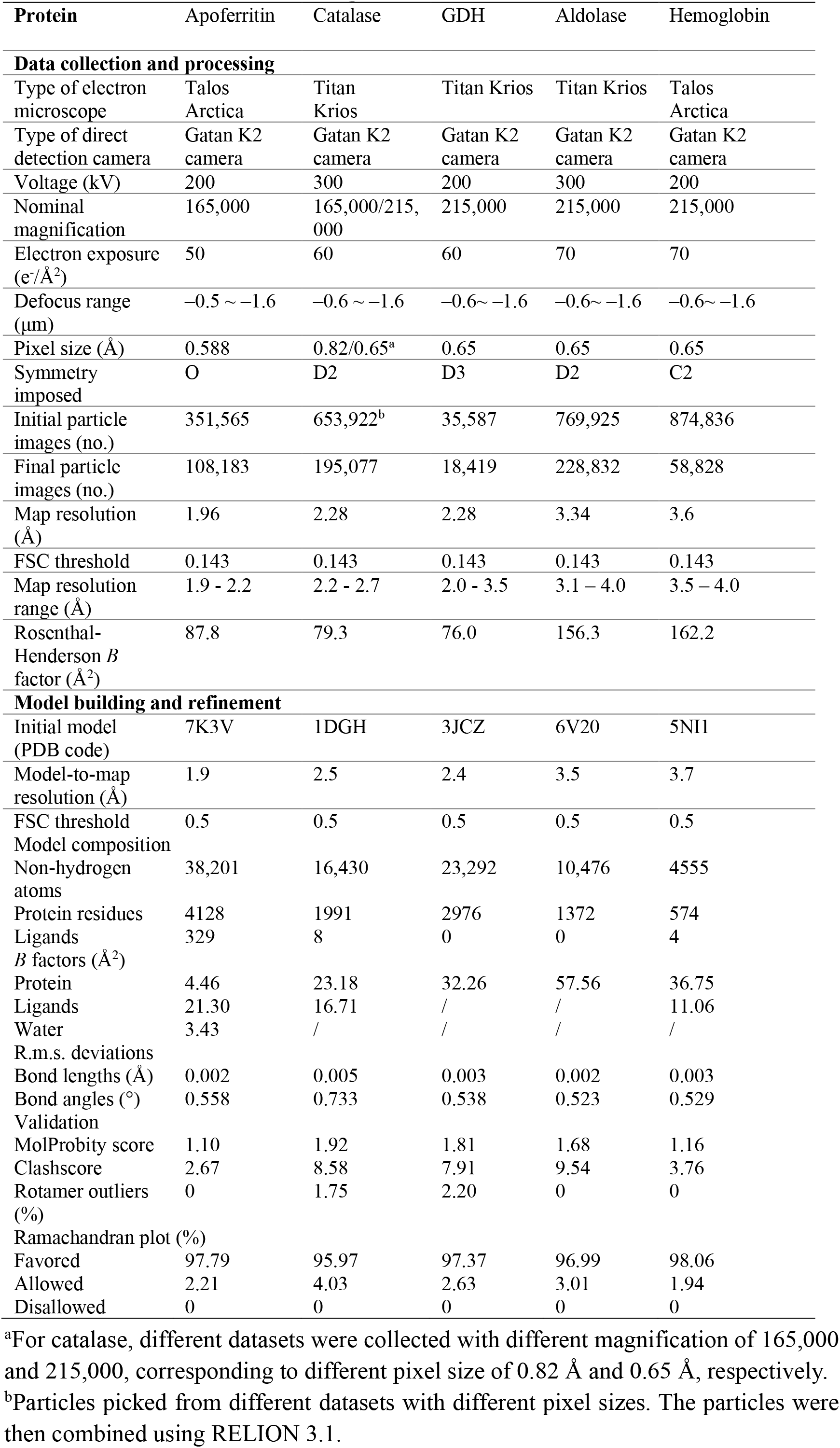
Statistics of cryo-EM data collection and processing, and model building, refinement and validation.

## Notes

### Competing Interest Statement

The authors have declared no competing interest.

